# Interleukin-4 signaling plays a major role in urogenital schistosomiasis-associated bladder pathogenesis

**DOI:** 10.1101/746552

**Authors:** Evaristus C. Mbanefo, Chi-Ling Fu, Christina P. Ho, Loc Le, Kenji Ishida, Michael H. Hsieh

## Abstract

IL-4 is crucial in many helminth infections, but its role in urogenital schistosomiasis, infection with *Schistosoma haematobium* worms, remains poorly understood due to a historical lack of animal models. The bladder pathology of urogenital schistosomiasis is caused by immune responses to eggs deposited in the bladder wall. A range of pathology occurs, including urothelial hyperplasia and cancer, but associated mechanisms and links to IL-4 are largely unknown. We modeled urogenital schistosomiasis by injecting the bladder walls of IL-4 receptor-alpha knockout(*Il4ra^−/−^*) and wild type mice with *S. haematobium* eggs. Readouts included bladder histology and *ex vivo* assessments of urothelial proliferation, cell cycle and ploidy status. We also quantified the effects of exogenous IL-4 on urothelial cell proliferation *in vitro*, including cell cycle status and phosphorylation patterns of major downstream regulators in the IL-4 signaling pathway. There was a significant decrease in the intensity of granulomatous responses to bladder-wall injected *S. haematobium* eggs in *Il4ra^−/−^* versus wild type mice. *S. haematobium* egg injection triggered significant urothelial proliferation, including evidence of urothelial hyperdiploidy and cell cycle skewing in wild type but not *Il4ra^−/−^* mice. Urothelial exposure to IL-4 *in vitro* led to cell cycle polarization and increased phosphorylation of AKT. Our results show IL-4 signaling is required for key pathogenic features of urogenital schistosomiasis, and that particular aspects of this signaling pathway may exert these effects directly on the urothelium. These findings point to potential mechanisms by which urogenital schistosomiasis promotes bladder carcinogenesis.

## Introduction

The pathogenesis due to schistosomiasis is chiefly a product of type-2 granulomatous immune response to the antigens secreted from the tissue-lodged parasite eggs [1]. The onset of egg deposition coincides with interleukin-4 (IL-4) production, the key inducer of Th2 response [2, 3]. The induction of Th2 response co-occur with the downregulation of Th1 response [4], and this immune phenotype is exploited by the parasite to complete their development to reproductive maturity [5], in addition to driving immuno-pathogenesis by orchestrating granulomatous response against egg antigens [6]. Indeed, in absence of this immune polarization, there is marked decline in tissue egg burden and egg-induced pathogenesis [7–10]. Egg secreted antigens induce release of cytokines to tightly regulate this otherwise lethal inflammatory response [11–13], allowing the parasite to survive for decades within the host; but unfortunately sets the stage for the characteristic fibrotic response. For intestinal schistosomiasis, hepatosplenomegaly and portal hypertension may result, or bladder cancer in the case of urogenital schistosomiasis.

The exact underlying mechanism by which schistosome eggs induce IL-4 production/release is still uncertain [2]. The major sources of IL-4 have also been shown to include other sources beyond the Th2 cells themselves [14]. Parasite egg crude extract has been shown to stimulate enormous IL-4 release from basophils [2, 15], mast cells and other non-T-cell, non-B-cell populations [16], which may indeed represent constitutive sources of IL-4, apart from the Th2 cells. Other Fcε receptor negative non-lymphocyte sources of IL-4 have also been reported [17, 18]. The most abundant IL-4 inducing egg antigen has since been identified and characterized from *S. mansoni* and *S. haematobium* [2, 15, 19, 20]. The interleuking-4 inducing principle from Schistosoma eggs (IPSE) binds to IgE on the surface of these non-T-cell sources, inducing massive IL-4 release that subsequently drive type-2 response [2, 15, 19, 20].

Although the role of IL-4 in driving schistosomiasis-induced pathogenesis is widely studied for hepato-splenic schistosomiasis, it remains to be fully shown whether similar mechanisms are present in urogenital schistosomiasis induced pathogenesis, which includes bladder carcinogenesis. This is mainly due to the lack of a tractable animal model and research reagents. We recently pioneered surgical introduction of eggs in the bladder walls as a model of urogenital schistosomiasis, reproducing most of the pathological changes associated with human *S. haematobium* infection in this intramural model [21–23]. Notwithstanding that urothelial cells have been shown to express IL-4 receptor [24, 25], there is paucity of studies on the role of IL-4 in urothelial changes in normal and disease states. Here, we examined the mechanistic role of IL-4 in the induction of bladder pathogenesis and carcinogenesis during urogenital schistosomiasis. We found that IL-4 receptor signaling is required for the recapitulation of the pathogenic features akin to human urogenital schistosomiasis. We further observed features consistent with oncogenesis and showed that the IL-4 effect on the urothelium driving bladder cancer is likely via signaling through the PI3K-Akt Pathway.

## Results

### Diminished bladder granuloma formation in egg-injected, IL-4 receptor-deficient mice

To understand whether the IL-4 receptor is required for the development of urogenital schistosomiasis associated bladder pathogenesis, we compared bladder granuloma formation in wild type BALB/c mice and IL-4 receptor knock out (IL-4Rα KO) mice (BALB/c background) following injection of *S. haematobium* eggs into the bladder wall. Wild type BALB/c mice and IL-4Rα KO mice were challenged with 3000 *S. haematobium* eggs by surgical intramural injection into the bladder wall. Ultrasonographic examination was performed 9 days post injection to evaluate the size of the granulomas. As shown in Figure 1, there was significant reduction in the size of bladder granulomas in IL-4Rα-deficient versus wild type mice (*p* = 0.0347) (Figure 1 and Figure S1).

**Figure 1.**
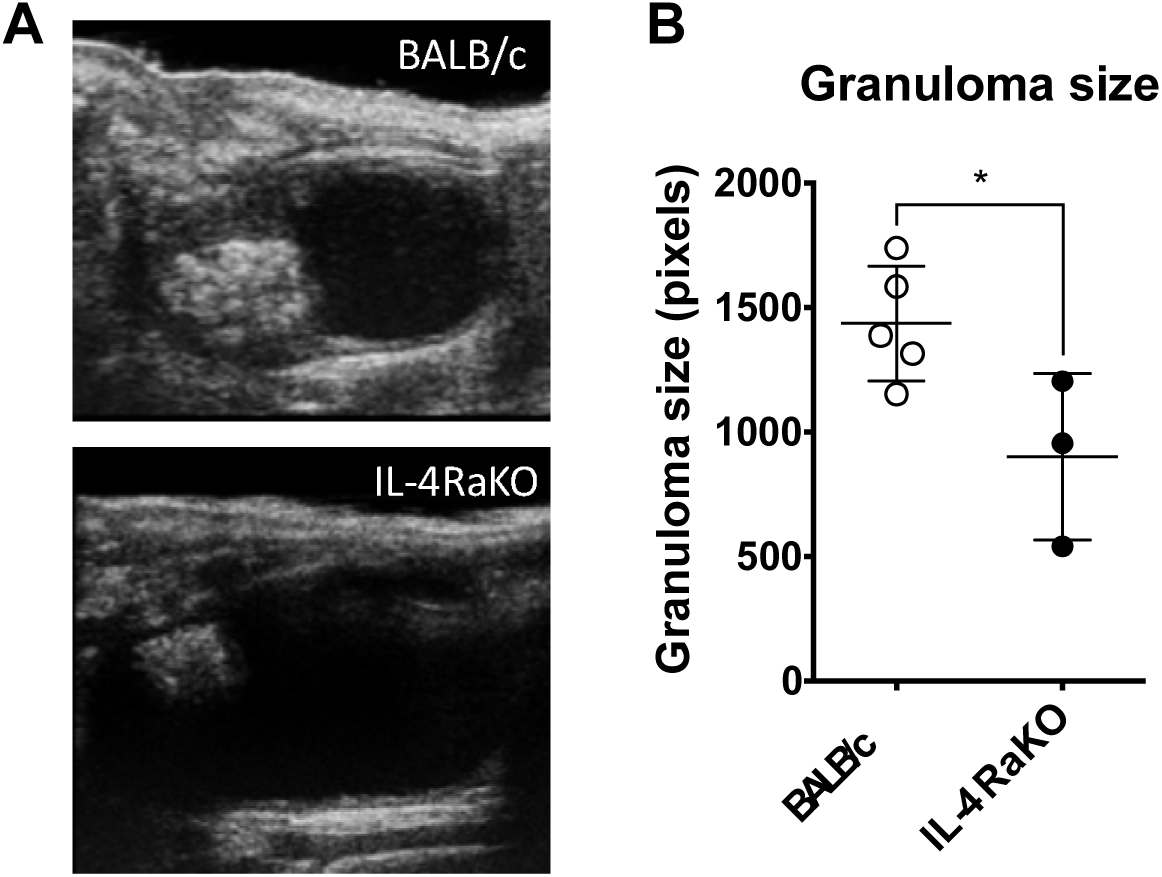
IL-4Rα KO mice demonstrate reduced egg-induced bladder granuloma size compared to wild type BALB/c mice. The bladders of BALB/c mice and IL-4Rα KO mice were intramurally injected with 3000 *S. haematobium* eggs. Granuloma size was evaluated 9 days post injection by ultrasonography. (A) Representative ultrasonographic images of the egg-injected bladders of BALB/c and IL-4RKO mice. (B) Measurements of sonographic granuloma size (also see Figure S1). The size of bladder granulomas in IL-4Rα-deficient mice was significantly reduced compared to wild type mice. Dot plots depict individual bladder granuloma size as analyzed using ImageJ. The midpoint of each dot plot shows the mean while the error bars represent one standard deviation.

We next conducted histologic examination of the egg-injected bladders. Low and high-power fields did not reveal any apparent differences in microscopic granuloma architecture (Figure 2). These data suggest that signaling via IL-4Rα helps determine the magnitude of the bladder granulomatous response during urogenital schistosomiasis but may be dispensable with regards to development of granuloma structure.

**Figure 2.**
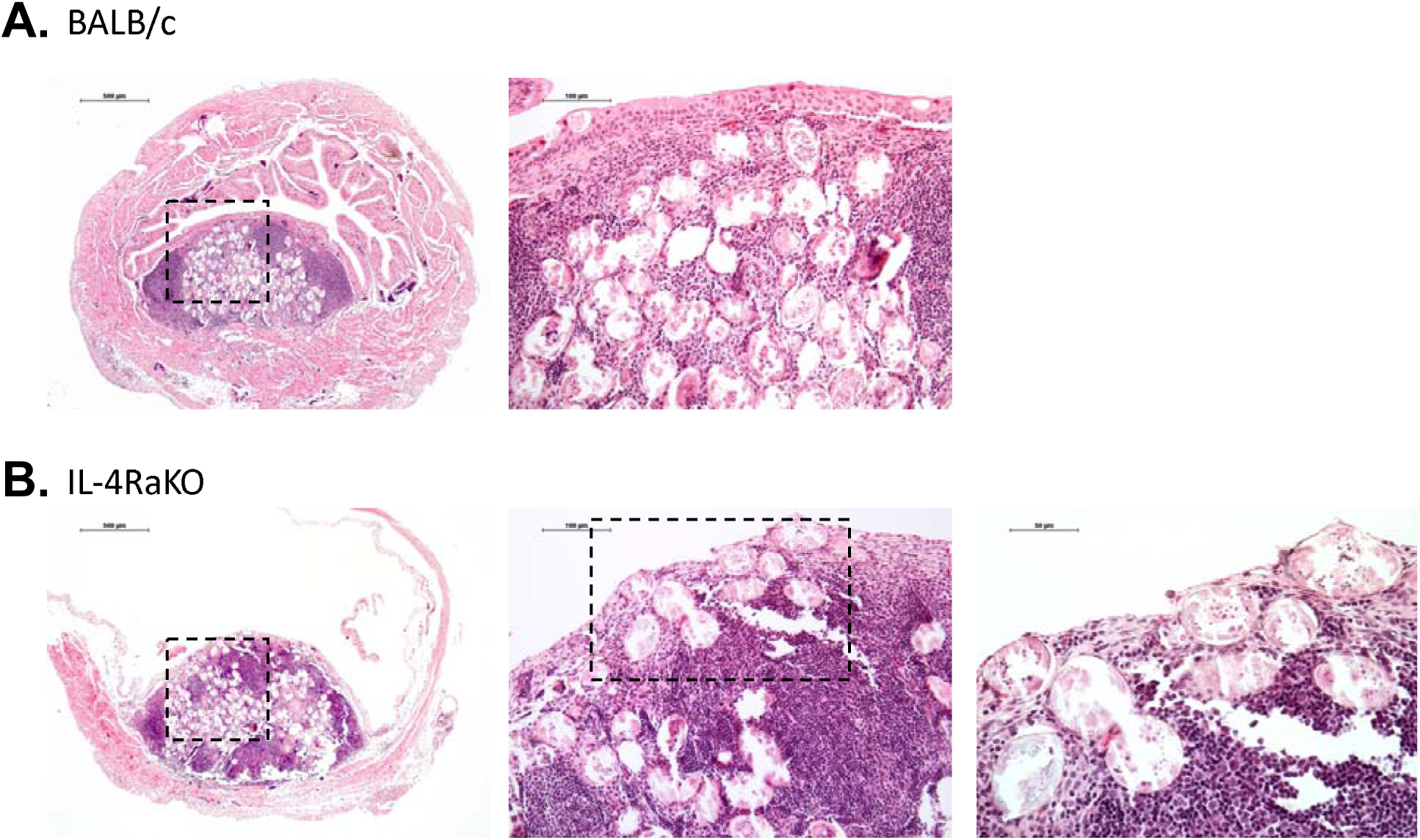
*S. haematobium* egg-induced bladder granuloma architecture is not IL-4 receptor-dependent. Wild type BALB/c mice (A) and IL-4Rα KO mice (B) were intramurally injected with 3000 *S. haematobium* eggs into the bladder wall. The bladders were excised 9 days post injection, fixed, dehydrated, embedded in paraffin, stained with hematoxylin and eosin, and sectioned. Representative low and high-power fields shown.

### *Schistosoma haematobium* eggs injection induce potentially pre-oncogenic cell cycle changes in urothelial cells

We next examined whether injection of *S. haematobium* eggs into the walls of a normal mouse bladder would produce potentially pre-oncogenic features in the urothelium, i.e., urothelial cell cycle polarization and development of hyperdiploidy. Mice were challenged with 3000 *S. haematobium* eggs by injection into the bladder wall. Bladders were excised, enzymatically processed and EpCAM^+^Uroplakin Ib^+^CD45^−^ urothelial cells identified via flow cytometry. While no apparent difference was observed in the proportion of cells at the G0/G1 phase of the cell cycle, there was a statistically significant increase in the proportion of urothelial cells at the S-phase in the group of mice receiving intramural egg injections as compared to groups receiving sham or no injections. In addition, there was significant increase in the proportion of urothelial cells exhibiting hyperdiploidy in the group of mice receiving bladder wall egg injections (Figure 3).

**Figure 3.**
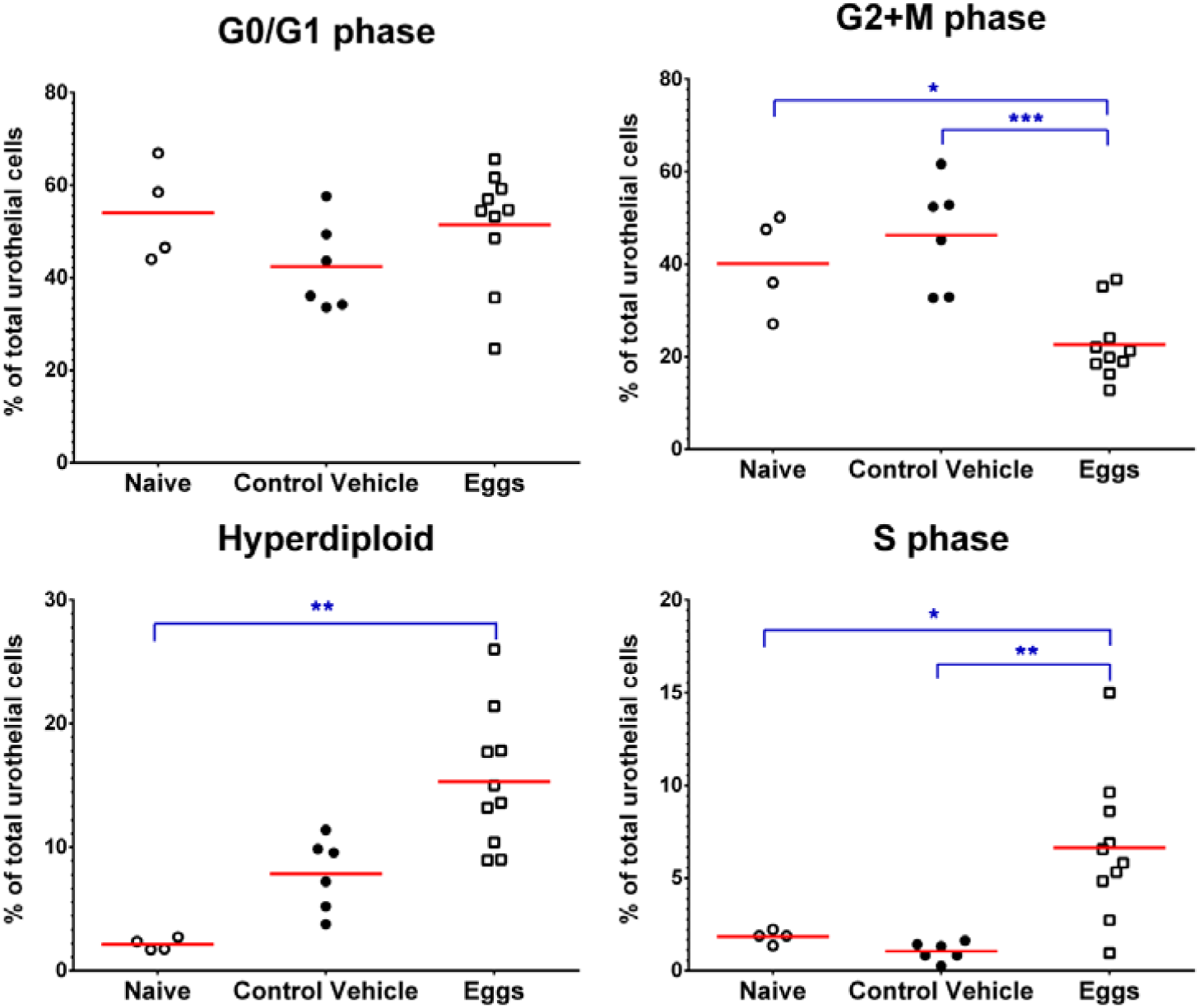
*Schistosoma haematobium* eggs drive potentially pre-oncogenic cell cycle and ploidy changes in the urothelium. Wild type BALB/c mice underwent bladder wall injection with 3000 *S. haematobium* eggs (isolated from infected hamster tissues) or vehicle (uninfected hamster tissue extract). Bladders were subjected to cell cycle analysis 4 weeks post-injection. Uroplakin 1b and EpCAM were used to isolate urothelial cells prior to cell cycle profiling (Figure S2). Compared to the vehicle-injected group, there was significant increase in the proportion of cells in the S-phase and decrease in G2+M phase cells following bladder wall egg injection. There was also significant increase in the proportion of cells showing hyperdiploidy in the egg-injected group. There was no difference in the proportion of cells in the G0/G1 phase. Red horizontal lines denote experimental group means.

### *Schistosoma haematobium* egg-induced urothelial cell cycle polarization is IL-4 receptor-dependent

Next, we sought to examine whether the observed *S. haematobium* egg-induced, potentially pre-oncogenic changes in urothelial cells were dependent on IL-4 receptor signaling. Wild type BALB/c and IL-4Rα KO mice were either challenged by bladder intramural injection with 3000 *S. haematobium* eggs or sham bladder wall injection of hamster liver and intestinal extract. At week 4 post-injection, all groups of mice, in addition to an unmanipulated naïve group of BALB/c mice, were administered BrdU to label actively proliferating cells. Twenty-four hours after BrdU administration, bladders were aseptically excised and subjected to flow cytometry to detect BrdU-labeled cells. As expected, there was a significant increase in the proportion of bladder cells in both S-phase and G2/M phase in wild type, egg injected vs. naïve mice (*p* = 0.0038 and *p* = 0.0026, respectively) and in wild type mice receiving a sham bladder wall injection (*p* = 0.0117 and *p* = 0.0003, respectively) (Figure 4). There was a statistically significant decrease in the proportion of S-phase cells in the bladders of egg-injected IL-4Rα KO vs. wild type mice (*p* = 0.0058, Figure 4). Similarly, the proportion of cells in the G2/M phase was significantly decreased in the egg-injected IL-4Rα KO mice (*p* = 0.0006), and sham treated IL-4Rα KO mice (*p* = 0.0009), as compared to egg-injected wild type mice. Interestingly, there was also a significant decrease in the proportion of S-phase cells in the egg-injected IL-4Rα KO mice (*p* = 0.0452) compared to the sham treated IL-4Rα KO mice (Figure 4B). Taken together, these results indicate *S. haematobium* eggs potently induce bladder cell cycle skewing in an IL-4Rα-dependent fashion.

**Figure 4.**
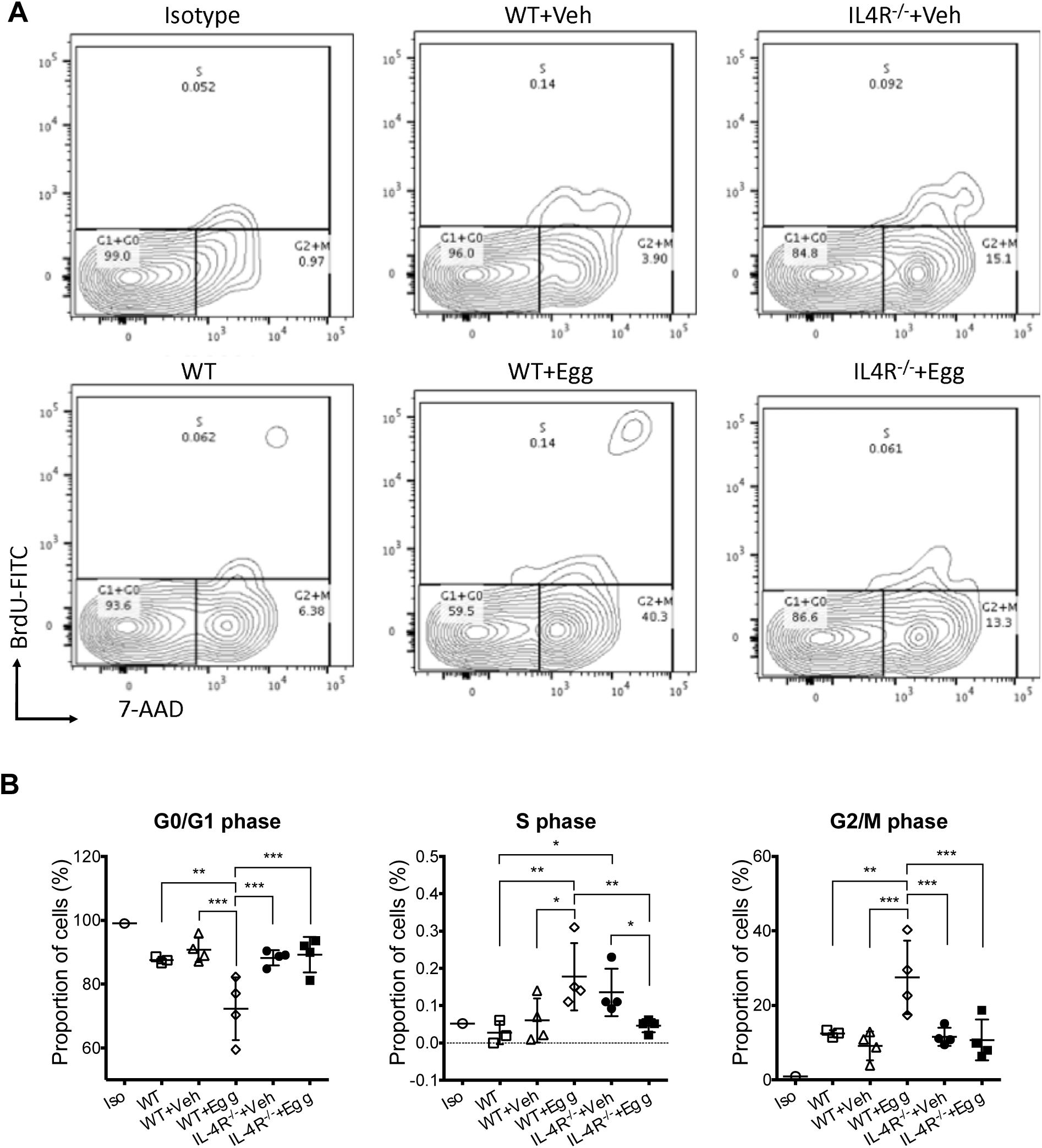
*Schistosoma haematobium* eggs drive cell cycle polarization in an IL-4 receptor-dependent manner. BALB/c mice or IL-4 receptor-deficient mice underwent bladder intramural injection with 3000 *S. haematobium* eggs (from infected hamster tissues) or vehicle (uninfected hamster tissue extract). (A) Bladders were subjected to BrdU staining and cell cycle analysis 4 weeks post-injection. (B) Compared to the egg-injected wild type group, in the egg-injected IL-4 receptor-deficient group, there was a significant decrease in the proportion of cells in the S-phase and G2+M phase, with a commensurate increase in the proportion of cells in the G0/G1 phase. The IL-4 receptor-deficient group exhibited cell cycle status profiles akin to the sham-injected group. Horizontal bars in each dot plot shows the mean, error bars represent one standard deviation.

### IL-4 receptor expression in the bladder is predominantly on the urothelium

Given the clear effects of IL-4 receptor signaling on *S. haematobium* egg-induced bladder pathogenesis, we next sought to determine whether IL-4 can act directly on urothelial cells. We compared the expression levels of IL-4Rα in the urothelium as compared to the detrusor smooth muscle cells (the other major cell type in the bladder besides the urothelial cell) using real-time quantitative PCR (Figure 5). We observed an approximately 2-fold increase in the expression of IL-4Rα in the urothelium as compared to the bladder detrusor.

**Figure 5.**
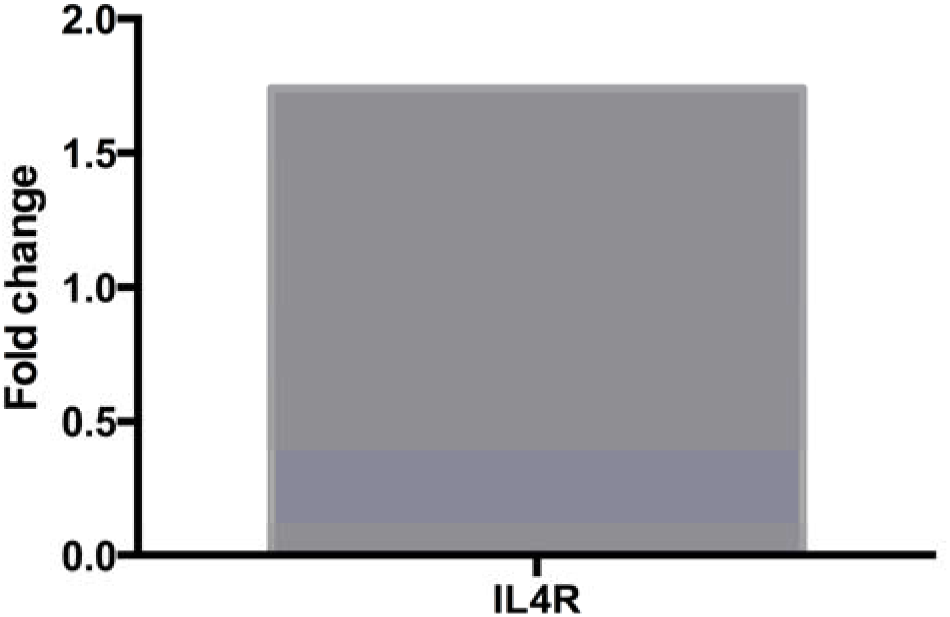
IL-4R expression in the bladder is mainly localized to the urothelium. Bladders were aseptically excised from mice followed by laser microdissection to isolate mouse bladder urothelium. Urothelial and detrusor cells were assayed for IL-4R expression using qPCR, and urothelial expression was calculated relative to detrusor expression.

### Exogenous IL-4 induces urothelial proliferation

To confirm that the observed, potentially pre-oncogenic changes in mouse urothelial cells were chiefly IL-4Rα-dependent and not due to other, unrelated effects of the parasite eggs, and to confirm our findings in a human model system, we performed *in vitro* assays using recombinant IL-4 and the human urothelial cell line HCV-29. HCV-29 cells were co-incubated with increasing concentrations of recombinant human IL-4, followed by assessment of cell proliferation and cell cycle changes. CFSE assays verified that IL-4 triggers urothelial proliferation (Figure 6). Cell cycle analysis revealed concentration-dependent increases in the proportion of cells in the G2/M phase at 0.1µg/ml (*p* < 0.0001) and 1µg/ml (*p* < 0.0001) of IL-4 (Figure 7). There was also a concomitant concentration-dependent decrease in the proportion of cells in the G0/G1 phase at 0.1µg/ml (*p* = 0.0008) and 1µg/ml (*p* < 0.0001) of IL-4.

**Figure 6.**
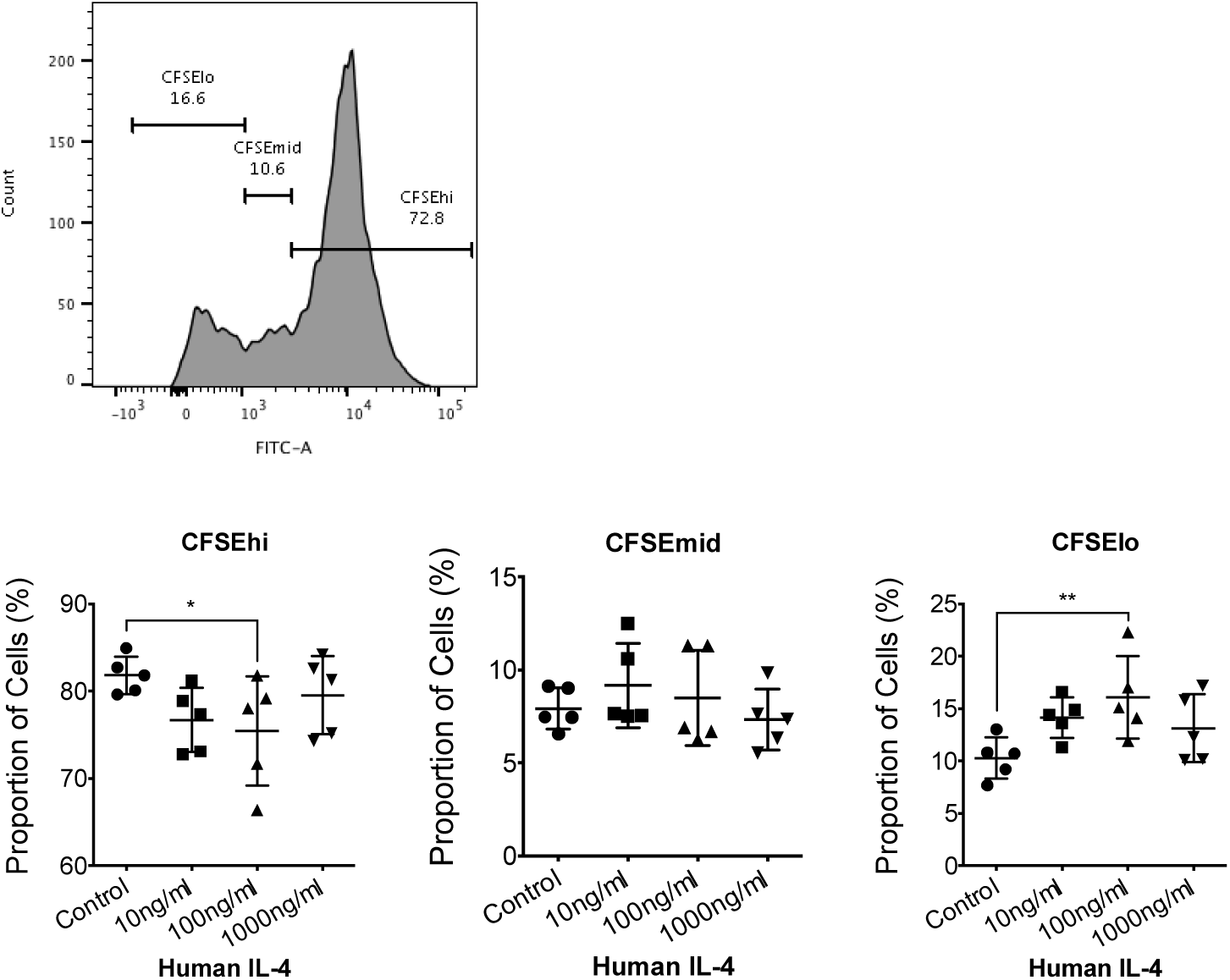
IL-4 directly drives urothelial proliferation. HCV-29 cells were stained with CFSE dye and then analyzed by flow cytometry to evaluate for cell proliferation as quantified by dye intensity. Upper left panel, example of gating strategy for CFSE intensity. G3 denotes cells that have not proliferated, G2 and G1 denotes cells that have proliferated and as a result feature lower levels of CFSE staining. Lower panels show proportions of cells as a function of CFSE staining intensity and IL-4 concentrations.

**Figure 7.**
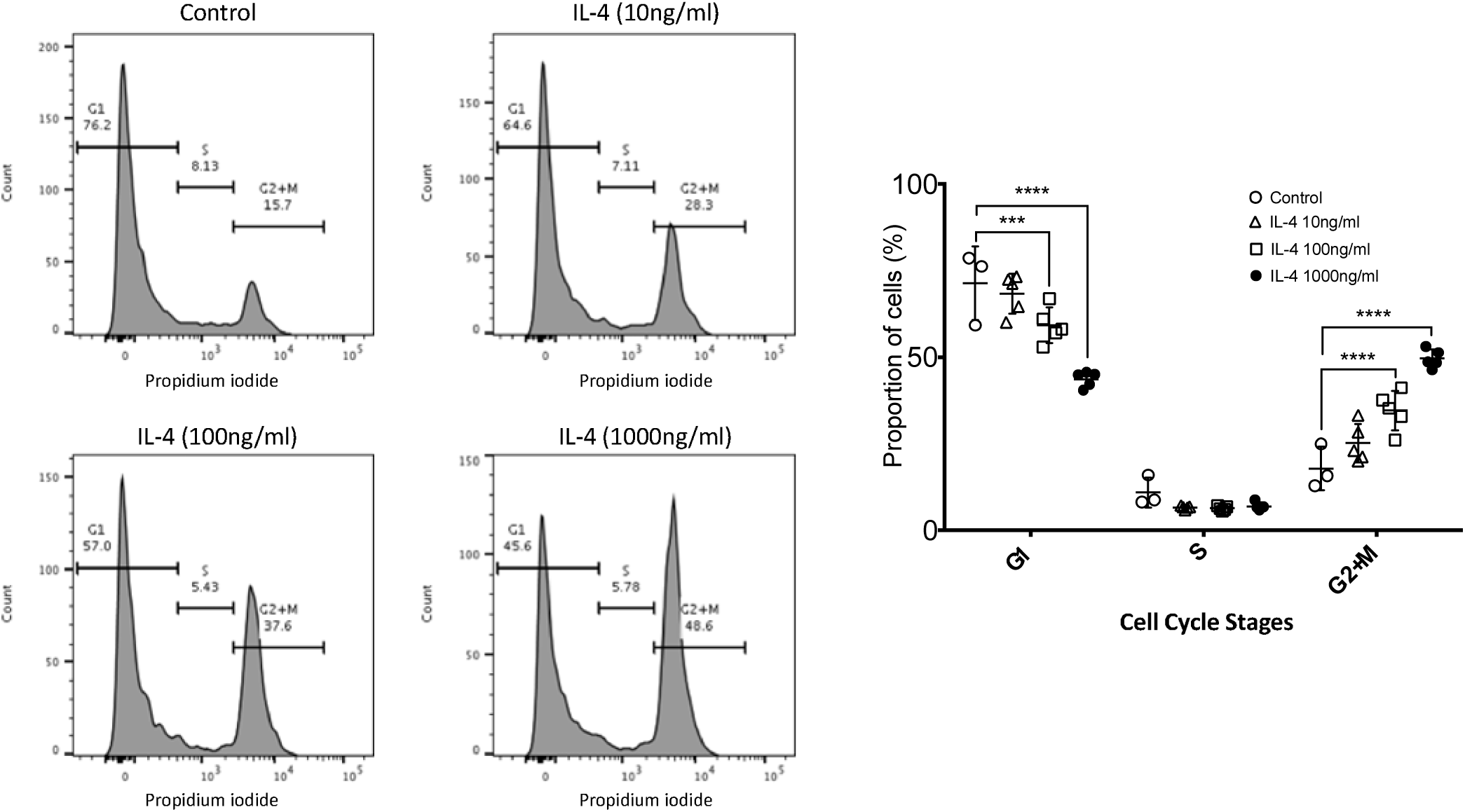
IL-4 directly induces urothelial cell cycle polarization *in vitro*. HCV-29 cells were co-incubated for 48 hours with increasing concentrations of recombinant IL-4, and then were subjected to cell cycle analysis. Propidium iodide staining of cellular DNA of fixed cells was used to evaluate cell cycle phases. Horizontal bars in the dot plots denote the mean and error bars represent one standard deviation.

### IL-4 drives urothelial proliferation via the IL-4R*α*/PI3K/AKT signaling pathway

IL-4 signaling progresses through two receptor complexes: heterodimers of the IL-4Rα and IL-2Rγc chain (type I receptor) and the IL-4Rα and IL-13Rα1 complex (type II receptor). These receptors trigger signaling via one or more of three cascades, viz: STAT6, PI3K/AKT and the ERK1/2 signaling pathway [26]. To ascertain the pathways through which IL-4 exerts its proliferative and cell cycle effects on urothelial cells, we used phospho-flow cytometry to ascertain the IL-4-induced phosphorylation status of relevant downstream proteins in MB49 mouse urothelial cells. Compared to urothelial cells incubated in the absence of recombinant mouse IL-4, and urothelial cells incubated with or without IL-4 but stained with sham antibody instead of phosphorylated AKT antibody, urothelial cells co-incubated with IL-4 and stained with phosphorylated AKT antibody showed a significant increase in the phosphorylation status of AKT protein (*p* < 0.0001) (Figure 8). We also found increased AKT phosphorylation in IL-4-exposed versus unexposed human urothelial HCV-29 cells (Figure S3). There was relatively less phosphorylation of STAT6 in HCV-29 urothelial cells, but this difference was nevertheless statistically significant compared to controls (p = 0.0077) (Figure S3). There was no change in the phosphorylation status of STAT6 and ERK1/2 (Figure S3). Moreover, bladder tissue from IL-4 receptor-deficient mice exhibited less AKT phosphorylation after *ex vivo* incubation with IL-4 compared to tissue from wild type mice (Figure S4). Taken together, these results show that IL-4 induces proliferative and cell cycle alterations in urothelial cells mainly via the PI3K/AKT signaling cascade.

**Figure 8.**
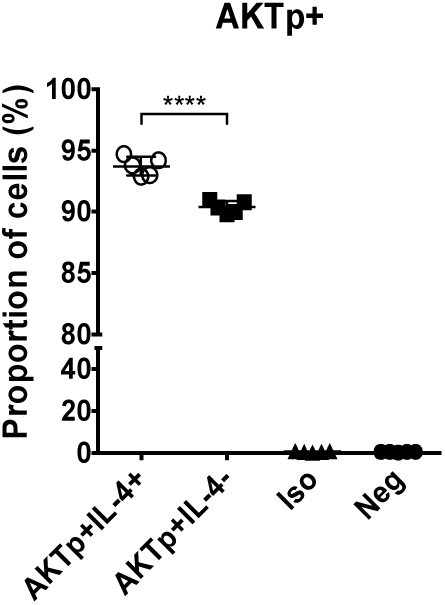
IL-4 directly increases AKT phosphorylation in urothelial cells. Mouse MB49 urothelial cells were incubated with recombinant IL-4 or no IL-4 and subjected to phosphor flow cytometry. “Iso”, isotype control antibody, “Neg”, no antibody staining.

## Discussion

IL-4 is arguably the classic cytokine associated with type 2 immune responses, including those directed against helminths (as reviewed by Webb and Tait-Wojno, among others [27]). Among human-specific helminths, schistosomes account for approximately 200 million infections worldwide, with the majority of these infections being urogenital and caused by *Schistosoma haematobium* [28]. Although IL-4 is also believed to be important in the bladder pathogenesis of urogenital schistosomiasis (chronic infection with *S. haematobium*), direct evidence of this hypothesized relationship has been scant. Much of the lack of supporting data has been due to difficulties with modeling urogenital schistosomiasis. We have previously shown that following micro-injection of purified *S. haematobium* eggs into the mouse bladder wall, most of the morphological features and pathological changes reminiscent of human urogenital schistosomiasis are reproduced in this tissue environment [1]. The response in this setting was rapid in onset and localized, including induction of IL-4. In other studies, we also showed that *S. haematobium* egg antigens induce IL-4 release from non-lymphocyte populations [20], mainly by activation via their surface Fcε receptors. Herein we sought to use our micro-injection technique to define the role of IL-4 in schistosomal bladder pathogenesis.

Infection with *S. haematobium* can result in a wide range of bladder pathogenesis, such as fibrosis, hyperplasia, dysplasia, squamous metaplasia, hematuria, and ultimately bladder cancer [29–32]. The link between *S. haematobium* infection and bladder cancer is so strong that the International Agency for Research on Cancer considers urogenital schistosomiasis as a class-1 carcinogen, “carcinogenic to humans” [33]. This parasitic infection is arguably one of the most important risk factor for bladder cancer globally, synergizing with other determinants like smoking, diet, host genetics, other environmental exposures and co-infection with other uropathogens [29]. Previously, we used p53 haplo-insufficient mice to demonstrate that loss of one allele of p53, in combination with *S. haematobium* egg injection, was sufficient to induce pre-carcinogenic lesions in the bladder [31]. We have also reported that massive shifts in DNA methylation occur in the mouse urothelium following bladder injection with *S. haematobium* eggs, and that these methylation changes drive urothelial proliferation, a necessary but not sufficient condition for bladder carcinogenesis [23]. Alterations in cell cycle regulation are associated with genomic instability, neoplasia and cancer progression, all of which are characteristic of urogenital schistosomiasis-associated bladder cancer [34]. We show here again that intramural injection of eggs into the bladder induces potentially pre-oncogenic cell cycle changes in urothelial cells, marked by increased S-phase populations and, interestingly, urothelial cell hyperploidy. Given that urothelial hyperplasia is triggered by *S. haematobium* eggs and is a feature of pre-neoplasia [31], this intramural egg injection model may be a step towards development of a tractable model of urogenital schistosomiasis-associated bladder cancer. Together, this body of work suggests that studying host responses to bladder wall injection with *S. haematobium* eggs may reveal important principles underlying schistosomal bladder pre-carcinogenesis.

One important principle underpinning schistosomal pathogenesis, independent of parasite species, is the importance of IL-4. For instance, the onset, development and progression of the granulomatous response and egg shedding during *Schistosoma japonicum* and *S. haematobium* infection, respectively, are regulated by the IL-4 signaling cascade [9, 35]. Signaling through the IL-4 receptor (IL-4Rα) also drives the chronic inflammatory response to the tissue-lodged eggs, which determines granuloma size, its cellular and matrix composition [10, 36], regulates the fibrotic repair response, and presumably subsequent carcinogenic sequelae. In fact, studies have demonstrated absent or significantly diminished granulomatous pathology in IL-4Rα-deficient mice compared to wild type and even IL-4-deficient mice [37]. This indicates that granuloma formation is not just dependent on the presence of IL-4, but rather also on the cascades downstream of IL-4Rα. Consistent with the foregoing, we used our intramural bladder wall egg injected mice model of urogenital schistosomiasis [21] to demonstrate significant decreases in granuloma size in IL-4Rα-deficient animals, indicating that signal transduction through IL-4Rα is required for the development of this central aspect of urogenital schistosomiasis induced immune pathogenesis.

We also present findings consistent with a role for IL-4Rα signaling in urogenital schistosomiasis-associated, potentially pre-oncogenic changes in the urothelium. *S. haematobium* egg-injected bladders from IL-4Rα-deficient mice showed significant decreases in the proportion of cells in both the S-phase and the G2/M-phases of the cell cycle as compared to egg-injected wild type mice and sham-injected IL-4Rα-deficient mice. Our emerging data from co-incubation of urothelial cell lines with *S. haematobium* orthologs of IPSE (Interleukin-4 inducing principle from *Schistosoma mansoni* eggs), show that IPSE and its IL-4-dependent effects may be one major driver of schistosomiasis-associated pre-oncogenesis (manuscript in preparation).

Prior work has pointed to tissue-specific roles for IL-4 receptor signaling in schistosomiasis. Namely, IL-4 receptor expression on alternatively activated macrophages and myeloid cells in general appears to drive specific aspects of *S. mansoni* pathology [38, 39]. IL-4 receptor expression by diverse cell types such as CD4+ and CD4-T cells [40, 41], smooth muscle cells [42], B cells [43], FoxP3+ regulatory T cells [44], and CD11c^+^ cells [45]. This corpus of research suggested to us that urothelial IL-4 receptor could be important in the bladder pathogenesis of urogenital schistosomiasis. Indeed, several studies have reported expression of IL-4 receptor in the bladder, presumably on the urothelium [24, 25]. After confirming that most IL-4 receptor expression in the bladder was on the urothelium, we exposed urothelial cells to exogenous IL-4 and found that IL-4 induces urothelial proliferation. Interestingly, this potentially pro-oncogenic effect is mainly via the IL-4Rα/PI3K/AKT signaling pathway and partly mediated through the STAT6 arm of the IL-4Rα cascade. This is particularly interesting because the PI3K/AKT and STAT6 arms of the IL-4Rα signaling cascade have been shown to be associated with promotion of cell proliferation and inhibition of apoptosis [26, 46, 47]. Indeed, Conticello *et al.* noted that IL-4 signaling enhanced urothelial and non-urothelial cancer cell resistance to CD95- and chemotherapy-triggered apoptosis [25]. Studies on several IL-4Rα-dependent mechanisms have also demonstrated pro-oncogenic effects of IL-4 signaling [46, 48], including promotion of cancer progression and metastasis in the bladder and other tissues [49, 50]. Conversely, strategies that block the IL-4Rα signaling cascade have been combined with cancer therapies to improve efficacy and reduce toxicity [26, 51]. Also, IL-4 and its fellow type 2 cytokine IL-13, as well as their cognate receptors, may have potential as biomarkers for tumor aggressiveness [47]. Venmar *et al.* showed that attenuation of IL-4Rα via the PI3K/AKT pathway reduces breast tumor cell survival and metastatic capacity [52]. All these studies are consistent with our observation that IL-4Rα signaling in the urothelium exerts a potentially pro-oncogenic effect, chiefly via the PI3K/AKT signaling pathway.

Although, to our knowledge, our observations regarding direct effects of IL-4 on the urothelium are the first to directly link type-2 immune responses to urogenital schistosomiasis and bladder pathogenesis, they may have relevance beyond *S. haematobium* infection. For instance, IL-4 may be important in other bladder-specific conditions, namely acute cystitis and overactive bladder [53, 54]. Future work focusing on urothelial IL-4 receptor signaling in various diseases may reveal novel diagnostic and therapeutic approaches and will promote our understanding of these conditions.

The work presented here has limitations worth noting. We have not ruled out an important role for leukocyte-urothelial interactions. Our *in-vivo* and *ex-vivo* urothelial observations in IL-4 receptor-deficient mice could have been confounded by the lessened granulomatous responses to eggs, which may have led to a dampened leukocyte-urothelial interface (whether mediated by cytokines and/or direct cell interactions). However, our *in-vitro* findings that IL-4 can mediate direct effects on urothelial cells indicate that it is possible that type-2-polarized immune responses may exert some of their effects on the bladder through IL-4-induced urothelial actions. Subsequent efforts will further focus on urothelial-specific IL-4 receptor signaling.

In conclusion, we have shown that IL-4 receptor signaling is required for the recapitulation of the pathogenetic features of urogenital schistosomiasis in the bladder. Namely, we showed that *S. haematobium* eggs induced urothelial proliferation, including evidence of urothelial hyperdiploidy. Interestingly, we further observed that these urothelial changes were dependent on IL-4 receptor signaling. The observation of features possibly consistent with pre-oncogenesis following urothelial exposure to IL-4 underline the potential importance of IL-4R signaling in pre-oncogenesis, and specifically urogenital schistosomiasis-associated bladder carcinogenesis. These IL-4-induced urothelial cell proliferation changes and potentially bladder carcinogenesis appear to involve the PI3K/AKT signaling cascade.

## Materials and Methods

### Ethics statement

Animal experiments reported here were carried out in accordance with relevant U.S. and international guidelines. Experimental protocols were reviewed and approved by the Institutional Animal Care and Use Committee (IACUC) of the Biomedical Research Institute (BRI), Rockville, Maryland, United States. These guidelines comply with the U.S. Public Health Service Policy on Humane Care and Use of Laboratory Animals.

### Mice, cells and reagents

Female BALB/c mice and IL-4Rα KO mice (*Il4ra^−/−^*, BALB/c genetic background) were obtained from Jackson Laboratory and were housed in a 12-hour light/dark cycle with dry mouse chow and water available *ad-libitum*. The human bladder epithelium (urothelium) cell line HCV-29 (a generous gift from Paul Brindley, George Washington University) was grown in T-75 tissue culture flasks in DMEM (Gibco) supplemented with 10% heat-inactivated FBS (Gibco) and 1× antibiotic-antimycotic (100 U/ml penicillin, 100 µg/ml streptomycin, 0.25 µg/ml Amphotericin B; Gibco) under 5% CO_2_ at 37 °C. Recombinant IL-4 was purchased from BioLegend (BioLegend, USA).

### Isolation of Schistosoma haematobium eggs

Male Golden Syrian LVG hamsters infected with *S. haematobium* were obtained from the National Institutes of Health-National Institute of Allergy and Infectious Diseases (NIH-NIAID) Schistosomiasis Resource Center at BRI. As previously described [21, 29], the hamsters were sacrificed around 16-18 weeks after exposure, the livers and intestines were excised, which were then processed in a blender and passed through sieves of pore sizes of 450, 180, 100, and 45 microns using cold 1.2% NaCl solution. The material from the final 45-micron sieve was transferred to and swirled in a glass petri dish and purified parasite eggs were collected from the center of the dish. Uninfected hamster livers and intestines were similarly processed to prepare control (vehicle) hamster extract.

### Mouse bladder wall injection

As described previously [21, 29, 55], mice were placed under systemic isoflurane anesthesia, their abdomens depilated and disinfected, and locally administered bupivicaine and buprenorphine. The bladder was exposed through a laparotomy, and 3,000 *S. haematobium* eggs in 50 µl of PBS were intramurally injected into the submucosal layer (bladder wall). Control mice were injected with control/vehicle hamster extract. The incision was closed with 4-0 polyglycolic acid suture and 4-0 silk suture, followed by local application of antibiotic ointment.

### Ultrasonographic imaging and histology of mouse bladders

A VisualSonics Vevo 770 high-resolution ultrasound micro-imaging system with an RMV 704 scanhead [40 MHz] (Small Animal Imaging Facility, Stanford Center for Innovation in In-Vivo Imaging) was used to transabdominally image mouse bladders [21]. For histology, bladder tissues were fixed in neutral buffered formalin, dehydrated in a series of ethanol incubations, embedded in paraffin, and stained with H&E [21].

### BrdU cell cycle assay

Mice were administered 1 mg of BrdU (FITC BrdU Flow Kit; BD) by intraperitoneal injection. After 24 hours, bladders were harvested, minced, incubated in tissue dissociation media (RPMI 1640, 10% heat-inactivated FBS, 15 mM HEPES, 1× antibiotic-antimycotic, 100 U/ml collagenase type III) [56] for 1 hour at 37°C with shaking, and passed through a 70-micron nylon cell strainer. The resulting single cell suspension underwent fixation, permeabilization, DNase digestion, anti-BrdU antibody-FITC labeling, and 7AAD labeling, according to BrdU kit instructions. Cells were analyzed by flow cytometry using a BD FACSCanto II equipment with acquisition run on BD FACS Diva software and analysis performed on FlowJo version 10 software.

### Urothelial-specific flow cytometry

Mouse bladders were excised, cut in half, and digested in 2.5 mg/ml of dispase in PBS for 1 hour at ambient temperature with shaking. Digested tissue was passed through 100-micron nylon cell strainers, minced, digested with 0.05% trypsin-EDTA at 37 degrees Celsius for 30 minutes, and added to the cell strainer pass-through material. The material was re-strained through another round of 100 micron nylon cell strainers. Cells were then incubated with anti-mouse CD16/CD32 in staining buffer to block non-specific Fc-mediated interactions, and then incubated with antibodies specific for EpCAM, uroplakin 1b, and CD45. For cell cycle analysis, cells were resuspended in DAPI solution (0.1% Triton X-100 + 4 micrograms/mL DAPI in 1X PBS buffer) for 37° Celsius for 1 hour with shaking in the dark. Cells were then washed and resuspended with staining buffer prior to data acquisition on a flow cytometer.

### Cell cycle analyses and CFSE assay

The human bladder epithelium (urothelium) cell line HCV-29 was grown in T-75 tissue culture flasks in complete DMEM media (Gibco) under 5% CO_2_ at 37 °C. For cell cycle assays, 10^5^ urothelial cells were co-incubated with recombinant human IL-4 at 0, 10, 100, or 1000 ng/ml concentrations. Following 48 hours of culture, the cells were fixed and stained with propidium iodide for cell cycle analysis. For CFSE assays to assess cell proliferation, cells were stained with the CFSE dye prior to stimulation with IL-4 and cultured for 48 hours. The CFSE dye was evaluated post-culture by flow cytometry using the FITC channel. The intensity of CFSE dye, which halves with each cell cycle, was used to track the generations of urothelial cells.

### Phosphorylation analysis of IL-4R signaling pathway

For phosphorylation analysis, cells were cultured as described above and subjected to intracellular staining using the Cytofix/Cytoperm fixation and permeabilization kit (BD, USA) according to manufacturer’s instructions. Fixed and permeabilized cells were then stained with antibodies against the phosphorylated forms of STAT6, ERK1/2 or AKT (ThermoFisher, USA). The stained cells were analyzed by flow cytometry using a BD FACSCanto II equipment run with BD FACS Diva software and analyzed on FlowJo software.

### Laser capture microdissection and real-time quantitative PCR

Laser microdissection was used to isolate urothelial cells or detrusor smooth muscle cells from the bladder, followed by real time PCR to measure IL-4R expression. Following cDNA synthesis using the iScript cDNA synthesis kit (Bio-Rad, California, USA), real time PCR was performed using the iTaq universal SYBR Green Supermix, following manufacturer’s protocols. The primers used for real time PCR are as follows: *IL-4R* (5’-TGGATCTGGGAGCATCAAGGT-3’ and 5’-TGGAAGTGCGGATGTAGTCAG-3’). Relative gene expression was then analyzed using the C_T_ method with fold expression using the formula 2^(-∆∆C^) with GAPDH as the internal reference and the amplification signal from the detrusor cells as the baseline.

### Statistics

Data analysis was performed using GraphPad Prism, version 6.00. One-way ANOVA was performed for comparison across groups and if significant, *post hoc* student *t*-tests was then used for pairwise comparisons after confirming a normal distribution of the data. Plots show individual replicates with horizontal bars denoting means and error bars denoting one standard deviation. Statistical significance was designated as *p* < 0.05. In the figures, * = *p* < 0.05; ** = *p* < 0.01; *** = *p* < 0.001; **** = *p* < 0.0001.

## Acknowledgements

We gratefully acknowledge our funding sources (the Margaret A. Stirewalt Endowment, NIDDK R01DK113504 and NIAID R56AI119168) and Jared Honeycutt for technical assistance.

## Author contributions

Designed research studies (MHH), conducted experiments (ECM, CLF, LL, CPH, KI), acquired data (ECM, CLF, LL, CPH, KI), analyzed data (ECM, CLF, CPH, LL, KI, MHH), provided reagents (MHH), and wrote the manuscript (ECM, KI, MHH).

---

**Figure S1.**
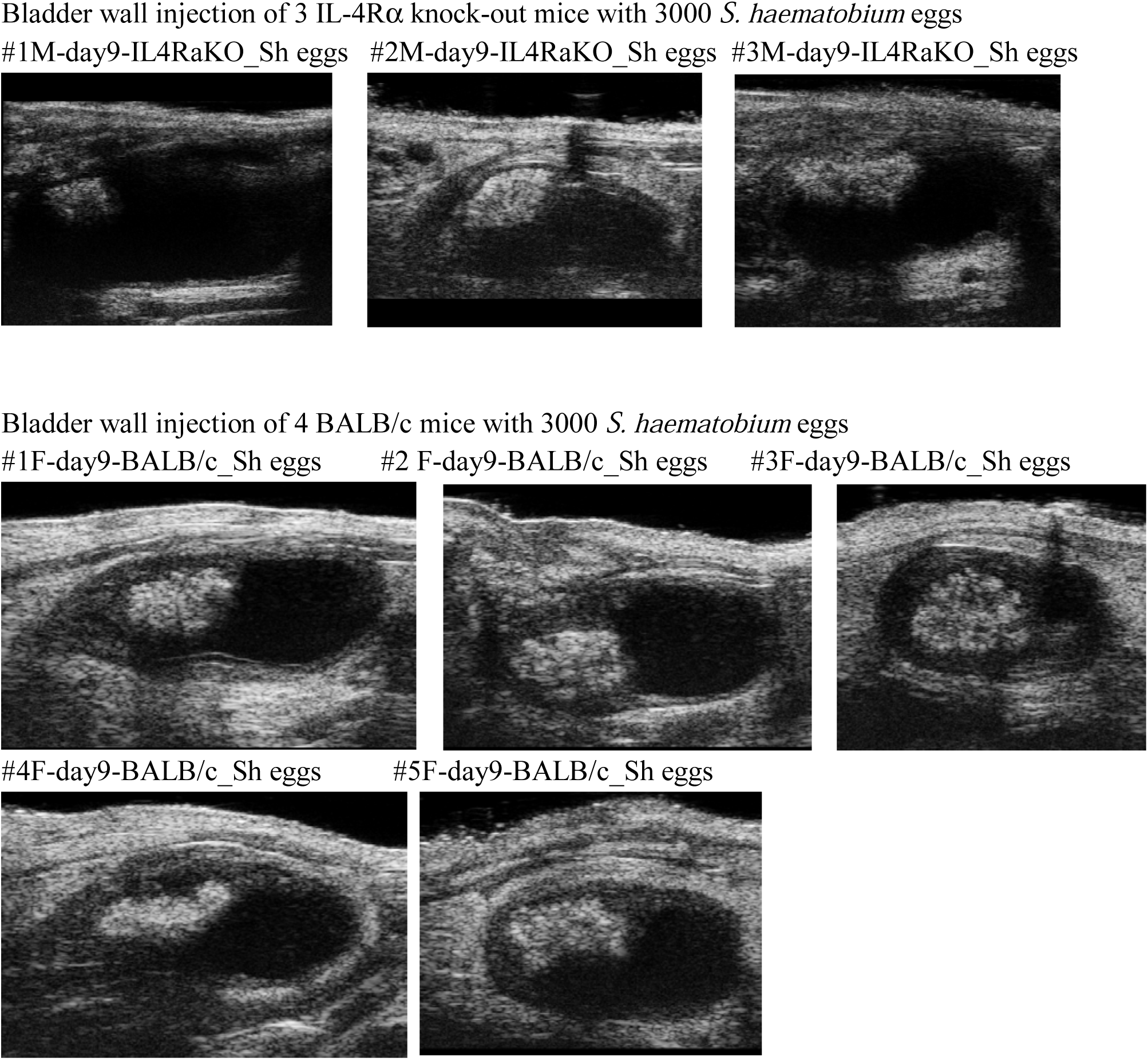
The size of egg-induced bladder granulomas is dependent on IL-4 receptor signaling.

**Figure S2.**
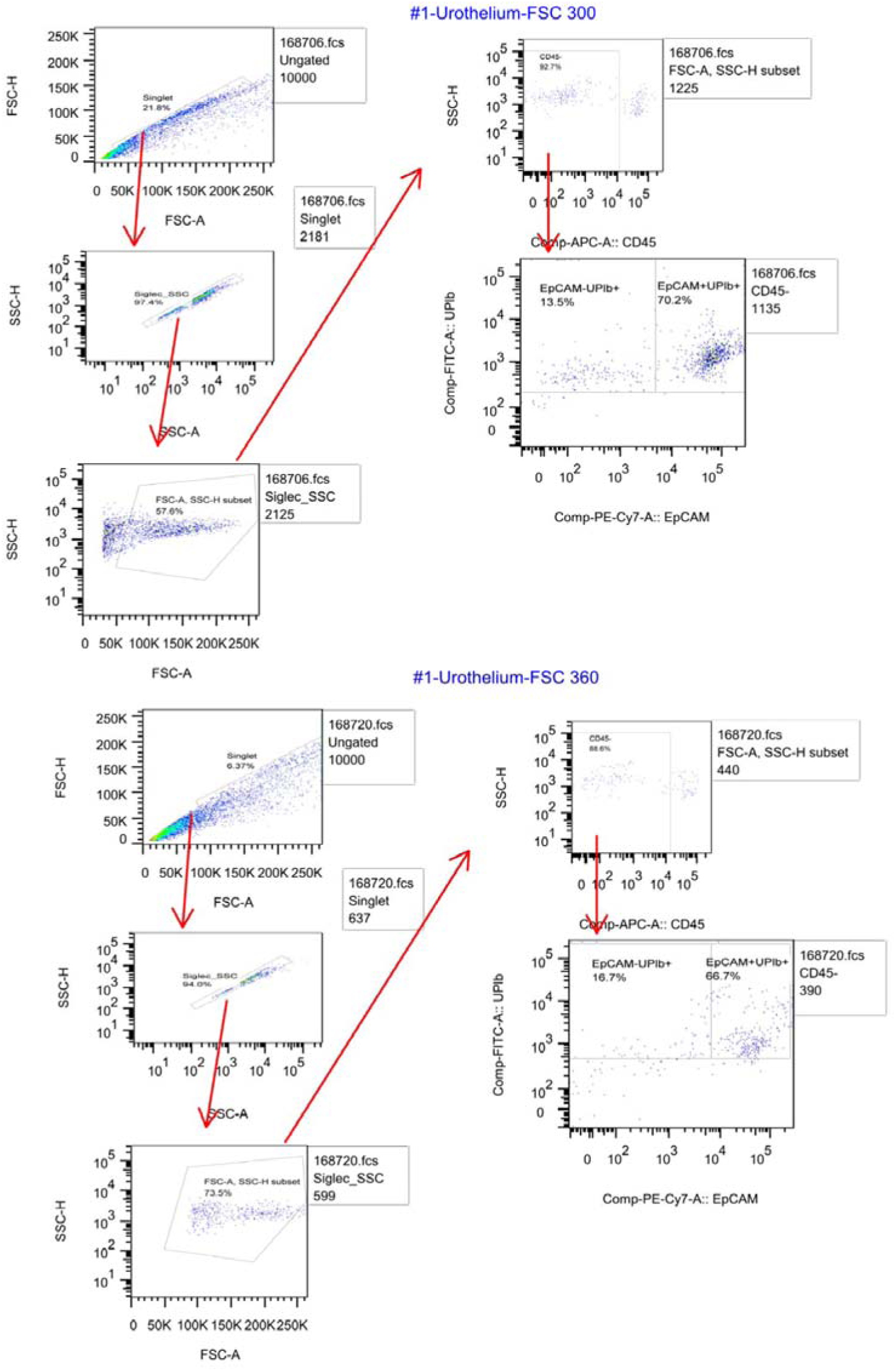
Gating strategy to enrich for EpCAM^+^Uroplakin Ib^+^CD45^−^ urothelial cells by flow cytometry.

**Figure S3.**
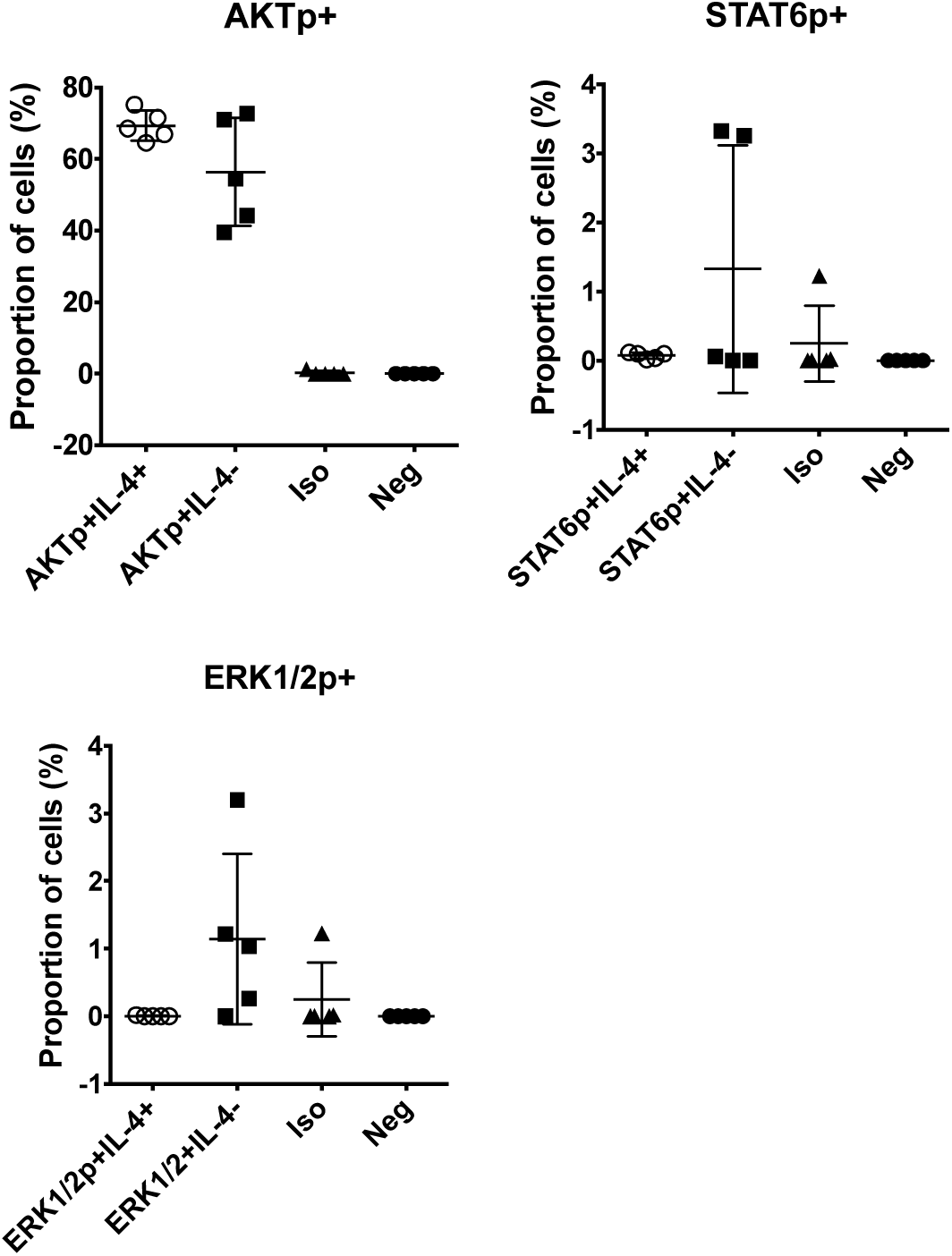
IL-4 drives urothelial proliferation via the IL-4 receptor/PI3K/AKT-signaling pathway. HCV-29 cells were co-incubated for 48 hours with IL-4, followed by intracellular staining of phosphorylated downstream signaling proteins in the IL-4R signaling pathway. Specifically, antibodies against phosphorylated forms of STAT6, AKT and ERK1/2 were used to assess the activation status of the three signaling cascades in the IL-4R signaling pathway. In the presence of IL-4, there was significant increase in the activation status of AKT compared to cells incubated without IL-4. The STAT6 pathway, in addition to ERK1/2, was also partially activated. Horizontal bars in each dot plot show the mean while the error bars represent one standard deviation. * = *p* < 0.05, ** = *p* < 0.01, *** = *p* < 0.001, **** = *p* < 0.0001.

**Figure S4.**
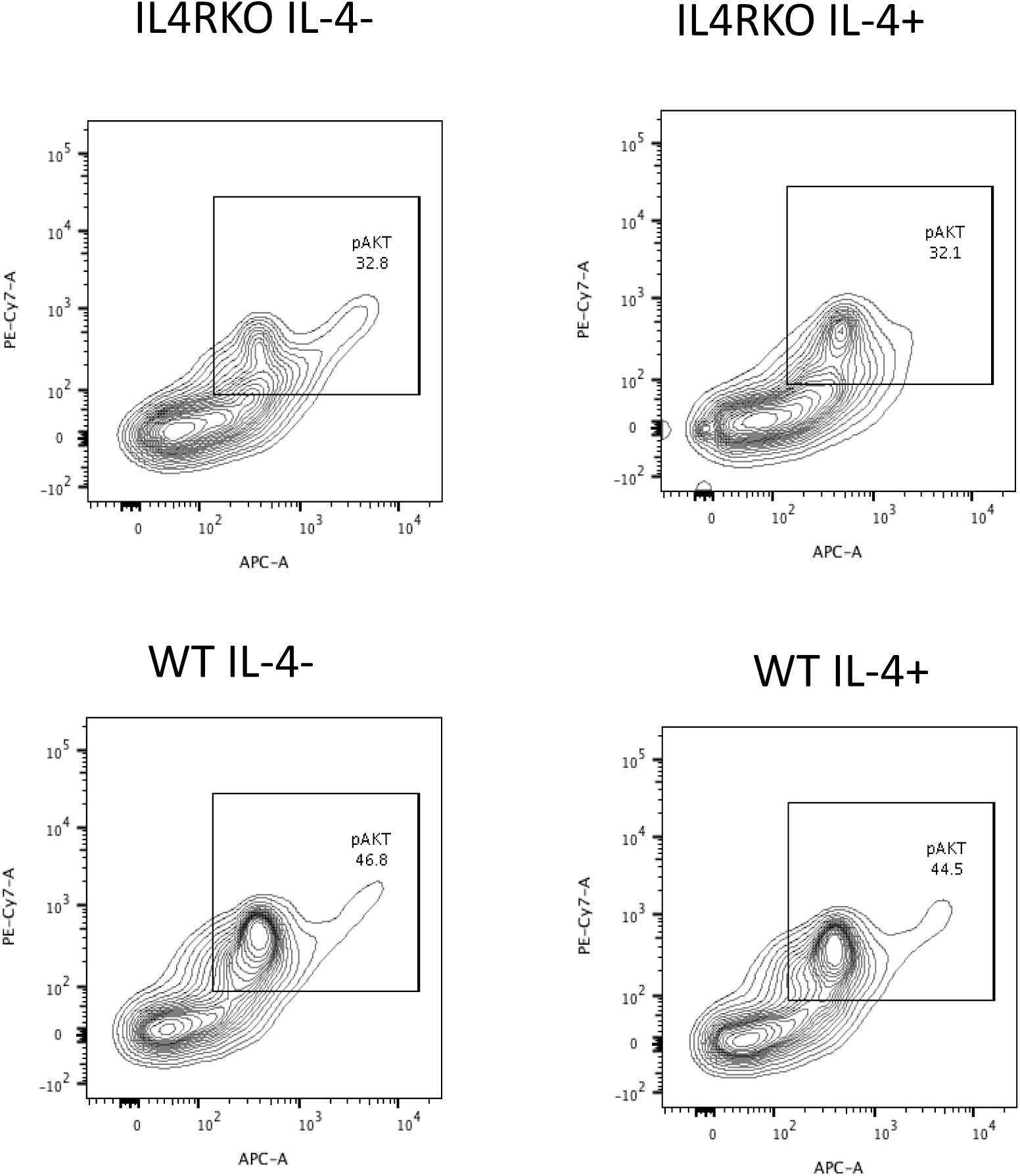
Primary mouse bladder tissue features increased AKT phosphorylation in response to incubation with IL-4 *in vitro* in an IL-4 receptor-dependent fashion. Bladders were excised from BALB/c or IL-4 receptor-deficient mice, incubated with recombinant IL-4 in culture, and subjected to Phospho flow cytometry using anti-pAKT antibodies.

